# Human motion data expansion from arbitrary sparse sensors with shallow recurrent decoders

**DOI:** 10.1101/2024.06.01.596487

**Authors:** Megan R. Ebers, Mackenzie Pitts, J. Nathan Kutz, Katherine M. Steele

**Affiliations:** Department of Applied Mathematics, University of Washington, Seattle, WA 98195; Department of Mechanical Engineering, University of Washington, Seattle, WA 98195; Department of Applied Mathematics and Electrical and Computer Engineering, University of Washington, Seattle, WA 98195

**Keywords:** gait analysis, machine learning, motion inference, sparse sensing, wearable sensors

## Abstract

Advances in deep learning and sparse sensing have emerged as powerful tools for monitoring human motion in natural environments. We develop a deep learning architecture, constructed from a shallow recurrent decoder network, that expands human motion data by mapping a limited (sparse) number of sensors to a comprehensive (dense) configuration, thereby inferring the motion of unmonitored body segments. Even with a single sensor, we reconstruct the comprehensive set of time series measurements, which are important for tracking and informing movement-related health and performance outcomes. Notably, this mapping leverages sensor time histories to inform the transformation from sparse to dense sensor configurations. We apply this mapping architecture to a variety of datasets, including controlled movement tasks, gait pattern exploration, and free-moving environments. Additionally, this mapping can be subject-specific (based on an individual’s unique data for deployment at home and in the community) or group-based (where data from a large group are used to learn a general movement model and predict outcomes for unknown subjects). By expanding our datasets to unmeasured or unavailable quantities, this work can impact clinical trials, robotic/device control, and human performance by improving the accuracy and availability of digital biomarker estimates.

## I. Introduction

HUMAN motion sensing and analysis is essential for monitoring disease progression, guiding rehabilitation, evaluating sports performance, and informing assistive device design. Biomechanics traditionally characterizes motion, such as gait, with digital biomarkers like kinematic, kinetic, and/or spatio-temporal parameters [3]. Certain biomechanical variables have been established as biomarkers that correlate with meaningful outcomes, such as knee adduction angle for Anterior Cruciate Ligament (ACL) injury or step width variability for fall risk, such as in aging populations [4]–[8]. In the US, where 1 in 7 individuals have a mobility disability and 1 in 2 adults live with a musculoskeletal condition [9], [10], monitoring motion ‘in the wild’ is vital for evaluating function and preserving mobility across the lifespan. For human motion to be observed in natural or uncontrolled environments, sensing devices must be portable, unobtrusive, reliable, and accurate. However, for sensing data to be meaningful, measurements must be converted to and contextualized as personalized outcomes, a challenge not yet overcome in natural environments [11].

While motion sensing is used across clinical, research, entertainment, and sports settings, the spectra of available technologies varies widely in practicality, accuracy, and expense; different environments dictate which sensor modalites can be used. Optical motion capture and force plates are considered the gold standard to comprehensively capture kinematics (motions) and kinetics (forces) [12]. However, these methods are highly specialized and require dedicated laboratories and careful calibration. This benefits only individuals near such clinics or who can afford the cost and time demands [13], [14]. Conversely, portable devices like inertial measurement units (IMUs), electromyography sensors (EMG), cameras, or insoles offer opportunities for out-of-lab motion sensing. However, these devices are less accurate than gold-standard optical motion capture to quantify human movement. For example, IMUs are widely used in wearable devices and have immense promise for motion sensing [15]–[19]. Questions of reliable placement by non-experts and error accumulation over extended use are major challenges, especially in natural environments [11], [20].

Machine learning has enabled researchers and scientists to push the boundaries of sensing and analysis of human motion using wearable sensors. Data-driven algorithms have proven to be powerful tools for extracting salient features and modeling complex relationships from the deluge of data collected from laboratory experiments, rehabilitation clinics, and wearable sensors. As a result, it has transformed human motion sensing and analysis for tasks like human activity recognition [21]–[26], markerless motion capture and pose estimation [27]–[34], fall detection [35]–[38], and sensor fusion [39]–[44], among others. Researchers have also investigated inference of human motion (*e.g*., kinematics) from reduced or sparse sets of sensors, in attempts to reduce the time and equipment burden. However, none result in comparable error to accurately monitor human movement compared to optical motion capture [33], [45]–[47]. Indeed, for highly-accurate motion sensing, laboratory-based technologies still prevail. A pipeline that enables comprehensive laboratory-level monitoring with wearables-level portability for natural motion remains elusive. To improve the accuracy and reduce sensor burden for natural environments, we can look to other fields facing sensor placement challenges, such as fluid dynamics. Researchers have developed state estimation techniques for accurately reconstructing a full set of states from a limited number of sensors [48]–[52]. In particular, a new deep learning method, the *SHallow REcurrent Decoder* (SHRED) network, reconstructs the full state space from limited measurements by learning a mapping from mobile sensor trajectories – which contain the time history of sensors – instead of only the current sensor value [1], [2]. SHRED uses (i) a recurrent network (*e.g*. LSTM) to learn a temporal latent space from the time histories of the limited measurements and (ii) a shallow decoder network to reconstruct the high-dimensional state space. This architecture – specifically because of its incorporation of the sensor’s time histories – outperforms other traditional (*e.g*., QR/POD [48]) or nonlinear (*e.g*., shallow decoders [52]) methods for state estimation, ensures robustness to measurement noise, and eliminates the need for optimally placed sensors [1], [2]. The shallow structure of the network requires significantly less data to train than typical deep learning architectures, which is ideal given the practical constraints of limited biomechanics data. SHRED also offers the capability of roll-outs, or future state prediction, through its recurrent neural network time-sequence encoding.

We theorized the SHRED algorithm could learn a mapping from a sparse set of wearable sensor measurements (*e.g*., triaxial accelerometer signals) to reconstruct dense sensor sets (*e.g*., full-body configuration of IMU signals) for both personalized and group analysis of human movement (Fig. 1). We hypothesized that incorporating time histories of sparse wearable sensor measurements would outperform linear and nonlinear techniques for data expansion to reconstruct human motion. We also aim to quantify the impact of training data (*e.g*. tasks with varying motion complexity) on expansion of motion data. We hypothesized that a SHRED model trained using datasets quantified as more complex would have lower data expansion error than a SHRED model trained on a dataset deemed less complex. To evaluate the performance of SHRED on human motion time series, we used four open-source datasets across a spectrum of movement tasks and measurement data, including treadmill walking and running, gait pattern exploration, and unconstrained activities like acting or freestyle dancing.

**Fig. 1.**
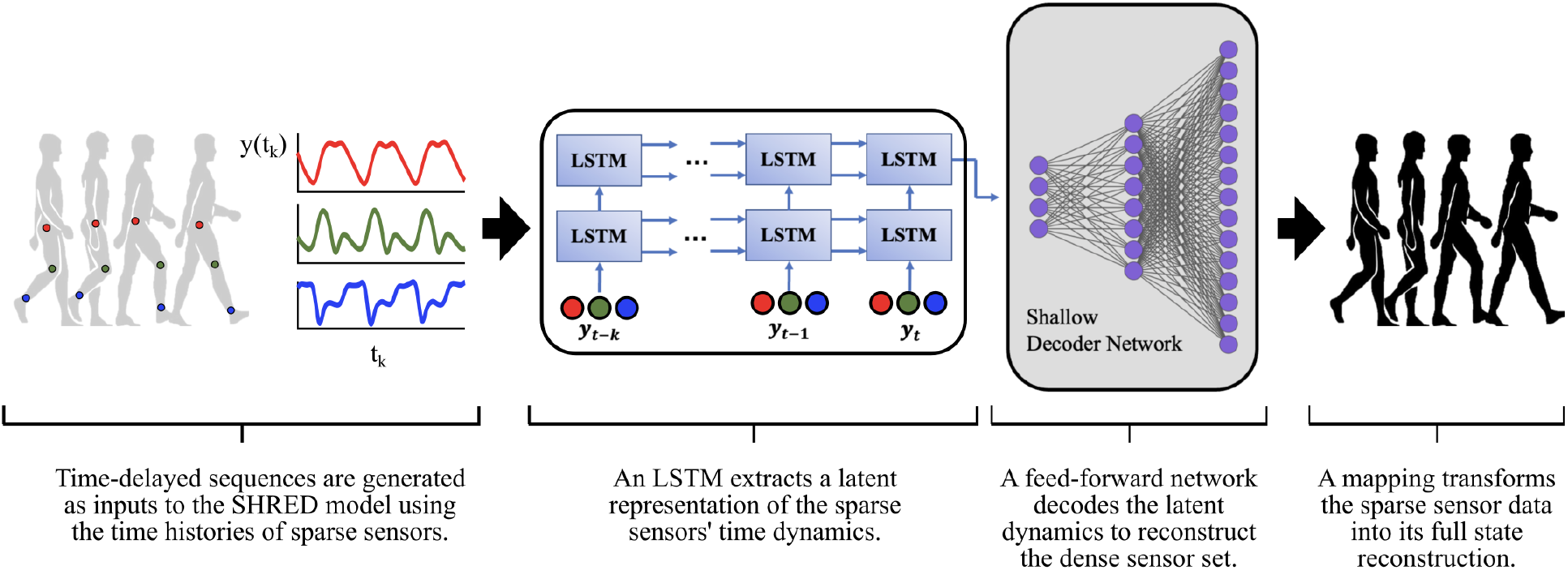
A deep learning and sparse sensing architecture enabling data expansion of human motion. For both personalized and group applications, we learn a mapping from a sparse set of sensor measurements to a dense sensor set using *SHallow REcurrent Decoder* (SHRED) networks [1], [2]. We leverage sensor time histories to learn a latent representation of human motion dynamics and reconstruct a comprehensive motion set in non-laboratory settings. We explore what data are needed to ensure robust data expansion.

## II. Methods

We organize our Methods in three parts: In Part A, we outline the mathematics of each modeling paradigm used to learn the mapping from sparse to dense sensor measurements, including *SHallow REcurrent Decoders* (SHRED), *Shallow Decoder Networks* (SDN), and *linear regression*. In Part B, we review the open-source datasets used to evaluate the utility of SHRED for data expansion. In Part C, we investigate the effect of motion complexity on motion inference.

We define data expansion as the transformation of sparse sensor measurements into a more comprehensive dataset, either through reconstruction or inference of the time series measurement(s). Reconstruction interpolates time series data for known data contexts, whereas inference predicts data for new subjects or tasks not previously encountered by the model.

### A. Overview of Models

We trained *SHallow REcurrent Decoders* (SHRED), *Shallow Decoder Networks* (SDN), and *linear regression* models to map sparse sensor measurements to dense sensor measurements. For each dataset and modeling paradigm, data were partitioned into a 60/20/20 training/validation/test set.

#### 1) Shallow recurrent decoder network

The *SHallow RE-current Decoder* (SHRED) is a deep learning architecture that maps the time history of sparse sensor measurements to its high-dimensional, spatio-temporal state. The architecture is expressed as:

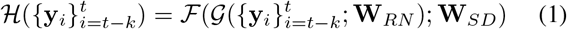

where **y**_*t*_ consists of measurements of the high-dimensional state **x**_*t*_, *ℱ* is a fully-connected, feed-forward neural network parameterized by weights **W**_*SD*_ and *𝒢* if an LSTM network parameterized by weights **W**_*RN*_. The SHRED architecture *ℋ* minimizes reconstruction loss,

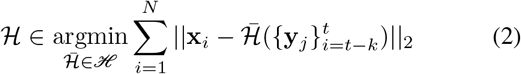

given a set of training states 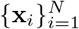 and corresponding measurements 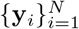 using the ADAM optimizer [53].

The data expansion accuracy is calculated as the magnitude of the error, normalized, between the ground truth and reconstructed/inferred time series:

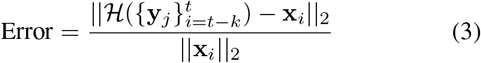

Due to SHRED’s reliance on time histories for state estimation and reconstruction, each dataset is truncated such that only the final *N −k* temporal snapshots are reconstructed/inferred, where *N* is the initial number of examples and *k* is the length of the utilized time history. In this manuscript, a time delay of *k* = 120 was used unless otherwise stated.

#### 2) Shallow decoder network

A shallow decoder network, which uses a fully-connect feed-forward network, can map sparse sensor measurements to a high-dimensional dataset. While this algorithm was part of the SHRED architecture, the shallow decoder network alone does not use the time history of measurements (*i.e*., no time-delay embedding nor LSTM):

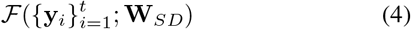

where **y**_*t*_ consists of measurements of the high-dimensional state **x**_*t*_ and *ℱ* is a fully-connected, feed-forward neural network parameterized by weights **W**_*SD*_.

Just as with the SHRED architecture, the shallow decoder network is trained to minimize reconstruction loss using the ADAM optimizer.

#### 3) Linear regression

Ordinary least squares regression fits a linear model with coefficients to minimized the residual sum of squares between the observed high-dimensional states in the test set with the states reconstructed by the linear approximation:

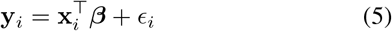

where 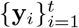 consists of measurements of the high-dimensional state 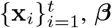 is the learned set of parameters, and 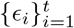 are unobserved random variables.

### B. Data Expansion of Human Motion

In this section, we implement each model for expanding human motion data and compare their performance. We began with a baseline task: nondisabled treadmill walking using two types of human motion data (marker-based and inertial). We further consider treadmill-based running to test data expansion of inertial sensors during a more dynamic task. Finally, we include additional motion complexity to understand the impact of gait pattern exploration (diverse motion) and environment exploration (movement constraints) on data expansion.

#### 1) Constrained movement: treadmill

We began with baseline treadmill tasks. If SHRED failed to reconstruct data that exhibits cyclic and symmetric characteristics, SHRED would reasonably fail to generalize to movement tasks with more complex features. We evaluated data reconstruction accuracy using error magnitude (normalized) for both reconstructing data from optical motion capture (joint angle kinematics) and wearable IMU sensors. We demonstrated two different types of mappings: (1) individual-specific mappings, which use each individual’s measurements to learn a unique mapping for personalized data expansion, and (2) group-based mappings, which uses all but one individual’s measurements to learn a group mapping and infers the held-out individual’s data, to test generalizability of a task’s mapping for unseen individuals. We compared variations of sparse sensor combinations to investigate the impact of sensor location and number.

##### Motion capture data

Rosenberg and colleagues collected motion data from 12 nondisabled adults (six female/six male; age = 23.9 ± 1.8 years; height = 1.69 ± 0.10 m; mass = 66.5 ± 11.7 kg) during treadmill walking at a self-selected speed (1.36 ± 0.11 m/s); data available: https://simtk.org/projects/ankleexopred [54]. Marker motion was recorded using a 10-camera optical motion capture system at 120 Hz (Qualisys AB, Gothenburg, SE), and joint kinematics were estimated using Inverse Kinematics in OpenSim 3.3 with a 19 degree-of-freedom skeletal model [55], [56].

Using joint kinematics, our goal was to learn mappings that reconstructed the full configuration of 18 kinematic states from a sparse set of measurements. For the individual condition, a mapping was learned for each of the 12 participants using the SHRED, SDN, and linear models. Four marker combinations were tested as map inputs: (1) three randomly chosen kinematic states (transverse-plane pelvis rotation, medio-lateral pelvis position, right hip adduction), (2) three purposefully chosen kinematic states (right hip flexion, right knee flexion, right ankle dorsiflexion), (3) one randomly chosen kinematic state (medio-lateral pelvis position), and (4) one purposefully chosen kinematic state (right ankle dorsiflexion). The purposefully chosen states were based on those most commonly used in clinical evaluations of walking [57], [58]. For the group condition, a mapping was learned using 11 of the 12 participants’ kinematic data, for a one-subject hold-out test, using the SHRED, SDN, and linear models. Two sparse sensor sets were tested: (1) three purposefully chosen kinematic states (right hip flexion, right knee flexion, right ankle dorsiflexion), and (2) one purposefully chosen kinematic state (right ankle dorsiflexion).

##### Inertial sensor data

Ingraham and colleagues collected inertial measurement unit (IMU) data from ten nondisabled adults (two female/eight male; age = 27.4 ± 4.5 yr; height: 1.76 ± 0.09 m; mass: 69.1 ± 9.9 kg; data available: https://doi.org/10.6084/m9.figshare.7473191) [59]. This team collected a variety of common steady-state and time-varying activities; for this example, we utilized their lower extremity and wrist IMU data from walking (0.9 m/s) and running (1.8, 2.2, and 2.7 m/s). Subjects wore four commercial triaxial accelerometers (Opal; APDM, Portland, OR) on their left and right lateral ankle, left hip, and the center of the chest. Each IMU measured acceleration (*m/s*^2^), angular velocity (*rad/s*), and magnetic field (*uT*) at 128 Hz in the x-, y-, and z-directions, totaling 36 distinct signals. Subjects also wore bilateral wristbands (E4; Empatica, Milan, Italy) containing triaxial accelerometers that recorded at 32 Hz.

With the IMU walking data, we evaluated two IMU configurations: (i) lower extremity IMUs only (left and right lateral ankle, left hip, chest center), and (ii) lower extremity + bilateral wrist IMUs. We trained SHRED, SDN, and linear models to expand sparse sensors to the full set of 36 lower extremity IMU signals for individual and group conditions.

##### Lower extremity IMU configuration during walking

Six IMU combinations were tested for their ability to reconstruct or infer 36 IMU signals: (1) triaxial left hip accelerations, (2) triaxial left ankle accelerations, (3) triaxial center chest accelerations, and (4) single-axis (x-direction) left hip accelerations, (5) single-axis (x-direction) left ankle accelerations, and (6) single-axis (x-direction) center chest accelerations.

##### Lower extremity + wrist IMU configuration during walking

Given that inertial sensors are ubiquitous in smartwatches, we also tested bilateral wrist IMUs as mapping inputs to reconstruct or infer the 36 lower extremity IMU signals. To ensure appropriate input-output mapping, the lower extremity IMU signals were downsampled from 128 Hz to 32 Hz to match the wrist IMU signal sampling rate.

##### Lower extremity IMU configuration during running

For each individual, we evaluated three scenarios for data expansion with SHRED during running: (i) single-speed (2.2 m/s), (ii) interpolation (train: 1.8, 2.7 m/s; test: 2.2 m/s), and (iii) extrapolation (train: 1.8, 2.2 m/s; test: 2.7 m/s). All models were trained using the right ankle IMU as the mapping input, as the distal tibia is a common sensor location for running studies [60]. One subject did not complete all the running trials, thus their data was excluded.

### C. Motion complexity and data expansion

#### 1) Gait pattern exploration

To investigate performance for more complex and diverse motion patterns, we used a dataset of gait pattern exploration. Spomer and colleagues collected marker motion data from 14 nondisabled adults (seven male/seven female; age: 24.1 ± 4.7 years; height: 1.7 ± 0.1 m; mass: 65.7 ± 20.1 kg) during treadmill walking at a self-selected speed (1.07 ± 0.13 m/s) [61]. Subjects performed one baseline walking trial and several exploratory trials, in which the “only imposed restrictions were that they must maintain forward-facing walking and take at least five consecutive strides in the pattern selected”. While this data was originally collected to understand whether individuals could voluntarily modulate their motor control complexity, these measurements provide a unique set of human motion data to quantify the effect of diverse movement on data expansion with SHRED. We trained individual-specific SHRED models for each baseline and exploration trial.

We quantified a task’s motion complexity and the effect of motion complexity on data expansion. If an individual explores more of their biomechanical space, will this provide a more robust signal that enables more accurate data expansion? We quantified motion complexity by calculating how much variance is explained by the first principal component (PC1) [62]–[64]; we performed PCA for every combination of tasks for each individual in the Spomer dataset. We hypothesized if a SHRED model is trained using the dataset quantified as being the most complex (*i.e*., the smallest PC1), it will have better reconstruction accuracy (*i.e*., smaller error) than a SHRED model trained on a less complex (*e.g*., random task combination) dataset. We identified the two individuals that demonstrated the smallest and largest change in motion complexity between trials. For these individuals, we (i) trained a SHRED model on their baseline data and predicted joint kinematics across all trials, (ii) trained a SHRED model on their baseline + most complex exploration trial and predicted joint kinematics during their most and least complex exploration trials, and (iii) trained a SHRED model on their baseline + least complex exploration trial and predicted joint kinematics during their most and least complex exploration trials. By understanding the effect of motion complexity – as a function of gait pattern exploration – on data expansion, we can evaluate how data used to train SHRED models impact robust data expansion across gait patterns.

#### 2) Environment exploration

We investigated environment exploration to understand the effect of movement constraints on data expansion with SHRED. By moving from a cyclic task like treadmill walking or running to tasks in a free-moving environment, we further probe the impact of motion complexity on data expansion. The TotalCapture dataset is a benchmark for computer vision tasks like pose estimation [65] and includes multi-view video, IMU, and marker (i.e., Vicon) labeling from 5 adults (four male/one female) each performing diverse tasks, including walking, acting, and freestyle dancing. We used a dataset from Van Wouwe and colleagues [66] that synthesized TotalCapture measurements using AddBiomechanics [67]. Similar to our gait pattern exploration approach, we quantified a task’s motion complexity during environment exploration by calculating how much variance is explained by the first principal component (PC1). We similarly hypothesized that if a SHRED model is trained using the dataset quantified as being the most complex (smallest PC1), it will have smaller data expansion error than a SHRED model trained on a less complex (*e.g*., random task combination) dataset.

## III. RESULTS

### A. Reconstruction accuracy across datasets

Shallow recurrent decoders (SHRED) successfully expanded human motion data by learning a mapping between a sparse set of measurements and the dense sensor configuration. For all datasets evaluated, SHRED models reconstructed motion data with lower error for both individual- and group-based scenarios than the other models. During treadmill walking, SHRED reconstructed joint kinematics with an average normalized error across individuals of 0.068 and 0.048 degrees for one and three sensors respectively (Table I). The comparison between SHRED and SDN reconstruction accuracy demonstrates the impact of time-delay embedding sensor time histories during model training, as seen in Fig. 2. For individual-specific mapping of joint kinematics during walking, SHRED was on average 4.0x and 12.8x more accurate than the SDN, for three and one sensor inputs respectively. For group-based mappings, SHRED was 3.3x and 9.8x more accurate than the SDN. While the SDN did learn a mapping between sparse and dense sensor configurations, linear regression resulted in high error for data expansion. SHRED outperformed SDN and linear regression for all datasets. We observed minimal impact of number or location of sensors on data expansion with SHRED (Table II). For example, when using IMU sensors during treadmill walking, we observed similar error for chest, pelvis, and ankle configurations, as well as trixaial and uniaxial accelerometer signals. Even for more dynamic tasks like treadmill running, IMU time series were reconstructed with low error regardless of number (one or three signals) or modality (accelerometer and/or gyroscope) (Table III).

**TABLE I.** Summary table comparing mapping methods for expanding joint kinematics during treadmill walking [54].

| Expanding joint kinematics ($^{\circ}$) during treadmill walking | | | | |
| --- | --- | --- | --- | --- |
| Type | Input Sensors | Error Magnitude, Normalized (± s.d.) | | |
|  |  | SHRED | SDN | Linear |
| Individual | 3 random | $0.048 \pm 0.026$ | $0.192 \pm 0.112$ | $0.359 \pm 0.261$ |
| | 3 purposeful | $0.055 \pm 0.032$ | $0.137 \pm 0.100$ | $0.439 \pm 0.345$ |
| | 1 random | $0.068 \pm 0.039$ | $0.871 \pm 0.542$ | $0.923 \pm 0.578$ |
| | 1 purposeful | $0.075 \pm 0.047$ | $0.516 \pm 0.303$ | $0.913 \pm 0.748$ |
| Group | 3 purposeful | $0.086 \pm 0.064$ | $0.281 \pm 0.170$ | $0.822 \pm 0.821$ |
| | 1 purposeful | $0.106 \pm 0.076$ | $1.041 \pm 0.839$ | $1.572 \pm 2.021$ |

**TABLE II.** IMU-based treadmill walking [59] comparing data expansion accuracy across mapping methods.

| Expanding IMU signals during treadmill walking |  |  |  |  |  |
| --- | --- | --- | --- | --- | --- |
| Type | | Input Sensors | error (± s.d.) | | |
|  |  |  | SHRED | SDN | Linear |
| Lower extremity | Individual | Triaxial Pelvic Accel. | 0.186 ± 0.133 | 0.541 ± 0.397 | 1.210 ± 1.204 |
| | | Triaxial Ankle Accel. | 0.243 ± 0.179 | 0.467 ± 0.352 | 1.713 ± 2.063 |
| | | Triaxial Chest Accel. | 0.179 ± 0.128 | 0.529 ± 0.381 | 1.245 ± 1.108 |
| | | Single-axis Pelvic Accel. (x-dir) | 0.437 ± 0.331 | 1.537 ± 1.415 | 2.757 ± 3.805 |
| | | Single-axis Ankle Accel. (x-dir) | 0.216 ± 0.152 | 1.070 ± 0.875 | 4.303 ± 7.923 |
| | | Single-axis Chest Accel. (x-dir) | 0.242 ± 0.173 | 1.271 ± 1.095 | 2.949 ± 4.037 |
| | Group | Triaxial Pelvic Accel. | 0.175 ± 0.134 | 2.680 ± 2.617 | 2.094 ± 2.334 |
| | | Triaxial Ankle Accel. | 0.178 ± 0.133 | 0.964 ± 0.656 | 2.801 ± 3.064 |
| | | Triaxial Chest Accel. | 0.178 ± 0.136 | 0.992 ± 0.681 | 1.971 ± 1.768 |
| | | Single-axis Pelvic Accel. (x-dir) | 0.197 ± 0.146 | 2.674 ± 2.580 | 4.255 ± 5.971 |
| | | Single-axis Ankle Accel. (x-dir) | 0.191 ± 0.141 | 1.885 ± 1.462 | 5.053 ± 7.523 |
| | | Single-axis Chest Accel. (x-dir) | 0.201 ± 0.150 | 2.198 ± 2.003 | 6.864 ± 20.003 |
| Wrist + lower extremity | Individual | Triaxial Bilateral Wrist Accel. | 0.213 ± 0.158 | 0.478 ± 0.348 | 1.062 ± 0.937 |
| | Group | Triaxial Bilateral Wrist Accel. | 0.266 ± 0.214 | 0.885 ± 0.637 | 3.316 ± 3.327 |

**TABLE III.** IMU-based treadmill running [59] evaluating single- and multi-speed data expansion accuracy.

| Expanding IMU signals during treadmill running |  |  |
| --- | --- | --- |
| Type | Input Sensors | error (± s.d.) |
| Single speed reconstruction<br>(2.2 m/s) | Uniaxial Ankle Accelerometer | 0.314 ± 0.177 |
| | Triaxial Ankle Accelerometer | 0.278 ± 0.169 |
| | Triaxial Ankle Gyroscope | 0.298 ± 0.167 |
| | Triaxial Ankle Accel. + Gyro. | 0.276 ± 0.168 |
| Interpolation<br>(train: 1.8, 2.7 m/s; test: 2.2 m.s) | Triaxial Ankle Accelerometer | 1.140 ± 0.298 |
| Extrapolation<br>(train: 1.8, 2.2 m/s; test: 2.7 m.s) | Triaxial Ankle Accelerometer | 1.255 ± 0.397 |

**Fig. 2.**
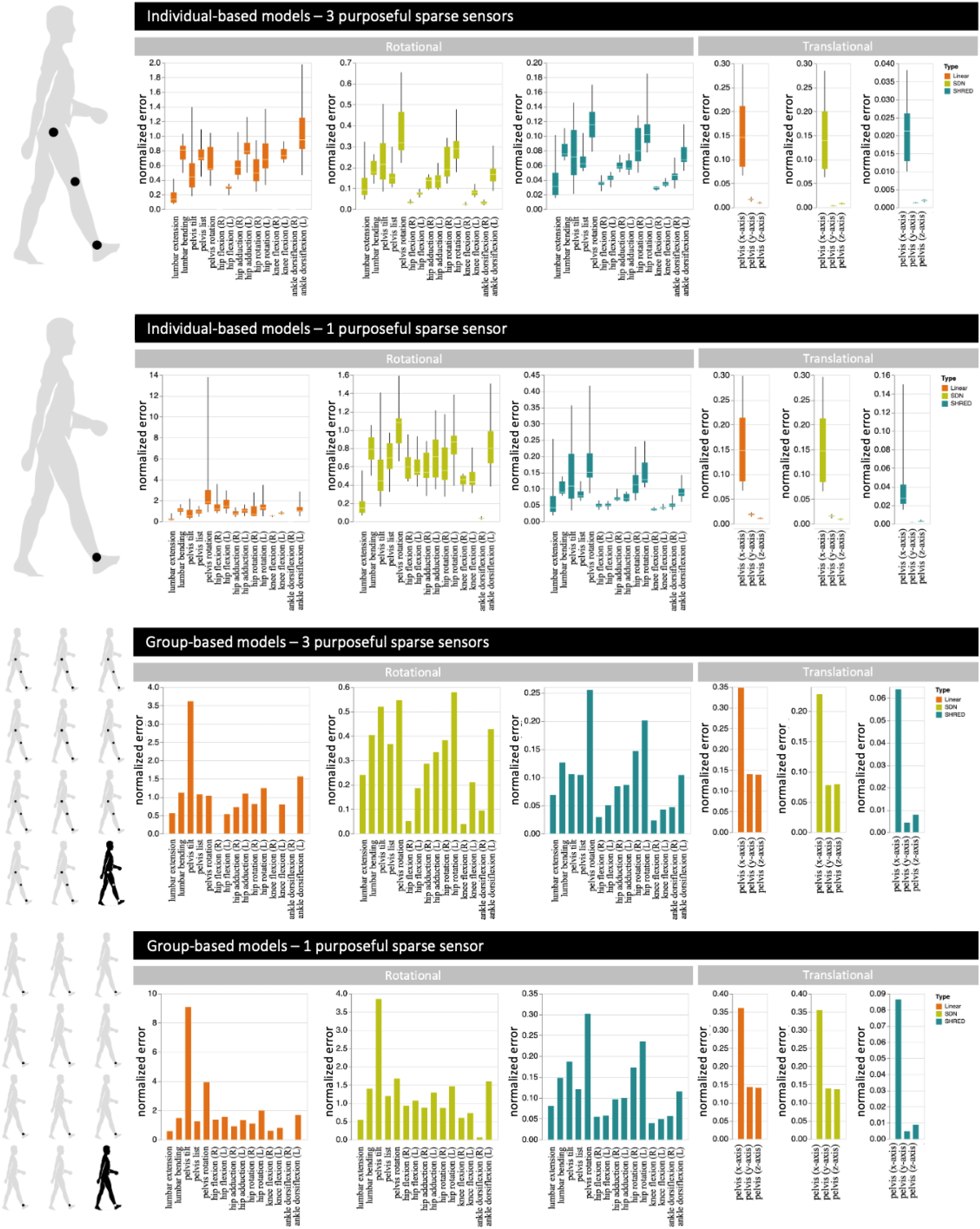
Shallow recurrent decoder (SHRED) models successfully expanded marker-based data during treadmill walking for both individual- and group-based scenarios [54]. Further, SHRED (blue) outperformed a shallow decoder network (SDN; green) and linear regression (orange). Performance was calculated using normalized error; box plots show the aggregate error for all individuals and group combinations, across kinematic states. Rotational [°] and translational [*m*] kinematic states are displayed separately. Note the independent y-axes and respective scales for each graph.

### B. Motion inference across datasets

SHRED also demonstrated lower errors for expanded data for new subjects and unseen tasks. Using SHRED, joint kinematics and IMU signals were inferred with low error for new subjects during treadmill walking (Tables I & II). For example, individual-based SHRED models inferred unseen kinematics from three purposely-placed sensors with average error of 0.055, with the average group-based SHRED error of 0.086 (SDN = 0.281, Linear = 0.822). Indeed, for inference during new tasks like unseen running speeds (error = 1.140 (interpolation) and 1.255 (extrapolation); Table III), diverse gait patterns (error = [0.8671, 1.1359]; Fig. 3), or unconstrained movements (error = [0.2825, 0.8922]; Fig. 4), SHRED inferred motion data from arbitrary sparse sensor configurations. The increased error and variability of data expansion with new tasks or subjects prompted investigation into the impact of training tasks on data expansion.

**Fig. 3.**
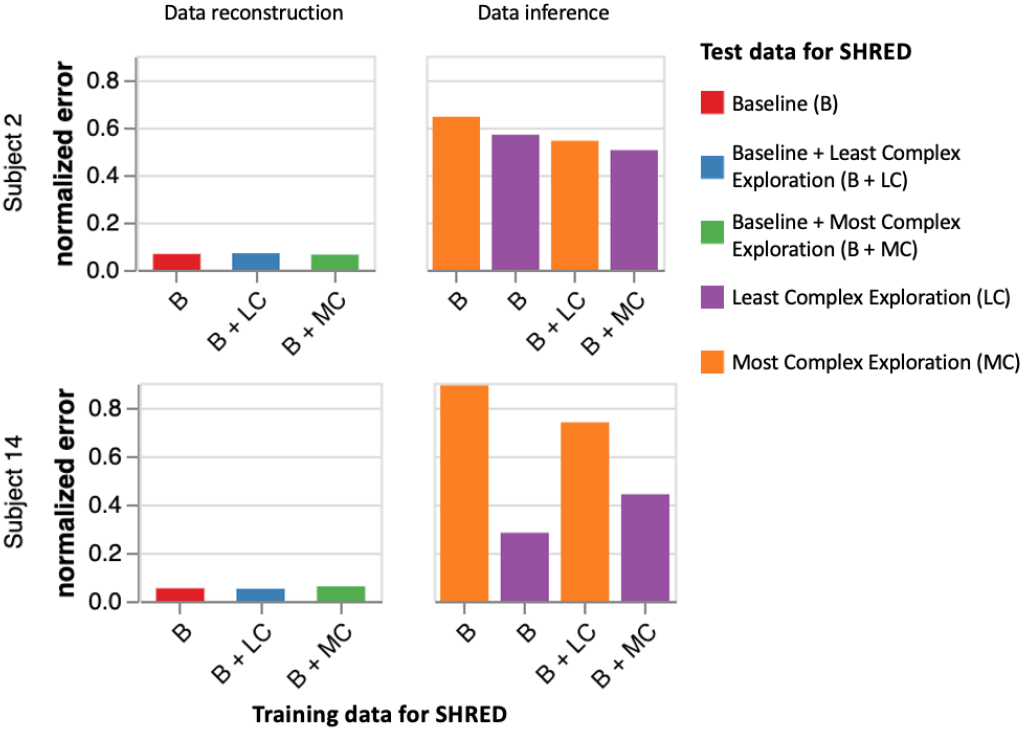
Increasing motion complexity improves data expansion accuracy predicting exploratory gait patterns [61]. We highlight data expansion results from two subjects; note: Subject 2 had the smallest change in motion complexity between baseline and exploration trials, while Subject 14 had the largest change in complexity.

**Fig. 4.**
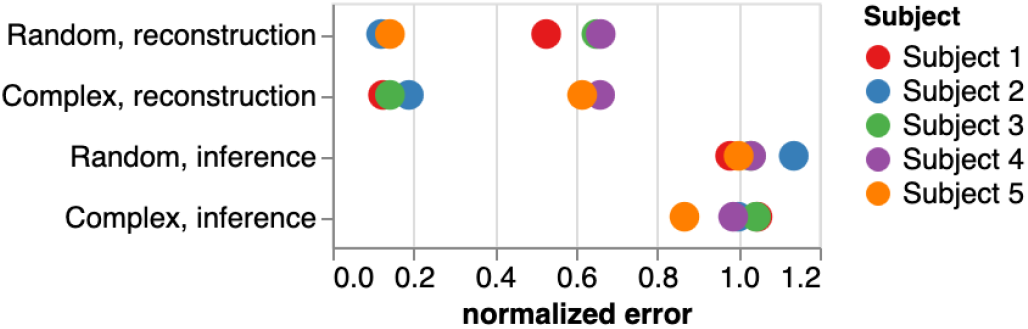
Increasing motion complexity had varied impact on data expansion accuracy with unconstrained movement tasks [65]. Individual-specific SHRED models were trained on either each individual’s most complex (PC1) task combinations or random tasks combinations. SHRED models expanded motion data by either reconstructing the full set of tasks or inferring an unseen task’s (*walking1*) data.

### C. Motion complexity and data expansion

#### 1) Gait pattern exploration

With gait pattern exploration, we observed that a change in motion complexity – as quantified by the first principal component (PC1) of joint kinematics – impacted motion inference. As seen in Fig. 3, we compared two subjects’ results; we chose these subjects to compare because Subject 2 (S2) showed the smallest changes in motion complexity across baseline and exploration trials (ΔPC1 of 0.0062), and Subject 14 (S14) showed the largest motion complexity changes (ΔPC1 of 0.1063). S2 and S14 had low error when reconstructing both baseline and exploration trials. We observed the largest impact of motion complexity on data expansion during motion inference. For example, S14 – who performed the most diverse gait patterns during their exploration trials – had a large difference in error between inferring their least complex (LC) and most complex (MC) gait explorations (LC=0.2825, MC=0.8922, Δ Error=0.6097); S2 – who performed the least diverse gait patterns during their exploration trials – had minimal changes in inference error across trials (LC=0.5693, MC=0.6447, Δ Error=0.0754).

#### 2) Environment exploration

With environment exploration, *i.e*., free movement in an unconstrained environment, we observed that a change in motion complexity had varied influence on motion inference. For all tasks, average reconstruction accuracy remained low (error=0.3457, for SHRED trained with complex tasks; error=0.4199, for SHRED trained with random tasks). On average, more complex training data reduced reconstruction accuracy, but the relationship between complexity and reconstruction accuracy varied between individuals (Fig. 4). For some subjects, like S1 and S3, greater complexity in model training resulted in improved reconstruction accuracy (S1: ΔPC1=0.1108, Δerror=0.4024; S3: ΔPC1=0.071, Δerror=0.5089). But for other subjects, like S2, increased training data complexity decreased reconstruction accuracy (S2: ΔPC1=0.0318 Δerror=-0.0688). We observed similar interpersonal variability with motion inference error.

## IV. Discussion

Our deep learning architecture expanded both individual- and group-based motion data by mapping a sparse set of sensor measurements to a dense configuration across movement tasks and environments. Shallow recurrent decoder networks consistently outperformed other linear and nonlinear mapping methods, highlighting the importance of leveraging the time history of sensor measurements in data expansion. However, as movement tasks become more complex, reconstruction accuracy decreased, which reinforces the importance of relevant training data and highlights the limits of motion inference.

This architecture’s expansion of motion data can advance efforts to improve the accuracy of digital biomarkers of movement-based health and performance outcomes. While wearable sensors show promise for personalized motion sensing, alone they struggle to capture critical biomarkers for health and performance [11], [68]–[71]. By using the methods proposed here to expand datasets from sparse wearable data to dense data akin to those obtained in a lab with individualized models, we may enable more continuous and less intrusive monitoring of movement-based biomarkers. Indeed, data expansion may enable more efficient and cost-effective natural monitoring, reducing the need for frequent visits to clinical or lab settings. To use these methods data collected during a motion capture session in a research or clinical gait analysis laboratory could be used to train a personalized SHRED model. This individualized model can then be paired with sparse sensing for future evaluations at home or in the lab. For example, in rehabilitation, progression is often challenging to track across sessions. An individual could have an early detailed assessment that could be used to train an individual SHRED model, such that future evaluations could use a single IMU to reconstruct and monitor specific biomarkers of interest. The time history of sensor measurements encodes a significant amount of information about human motion. It has been established that human gait can be represented as a nonlinear dynamical system, which inherently assumes that past behavior influences future behavior [72]–[75]. The shallow recurrent decoder architecture exploits this inductive bias by learning the temporal and spatial relationships between sparse and dense sensor configurations. Indeed, our results demonstrate that the time histories of a sparse sensor set are important for reconstructing unmeasured motion, using as few as a single IMU sensor. By comparing SHRED to shallow decoder network that do not utilize LSTM, to capture time histories, we demonstrated a distinct improvement in data expansion accuracy across sparse sensor configurations, number of sensors, sensor modalities, and both individual- and group-based mappings.

Previous approaches that successfully infer human motion, such as DiffusionPoser [4], utilized a combination of whole-body pose and IMU information from large group-based datasets. DiffusionPoser successfully reconstructs human motion from arbitrary IMU configurations with as few as two sensors. However, the neural network underlying diffusion models does not leverage sensor time histories, which may further improve motion inference. Additionally, incorporating skeletal representations to reconstruct motion may be inappropriate for some individuals, such as those with bony deformities, potentially limiting utility and interpretability in clinical populations [76]–[78]. Our motion inference approach is purely data-driven and highly personalizable. We utilized techniques rooted in dynamical systems, differential equations, and time-delay embedding [79]–[82]. Such techniques allowed us to uncover – rather than prescribe – structure underlying human motion from sensor measurements, enabling personalization while avoiding explicit physiological assumptions.

Motion complexity impacts motion inference. We hypothesized that a SHRED model trained using datasets quantified as more complex would have lower data expansion error than a SHRED model trained on a dataset deemed less complex. Across all tasks and for both individual- and group-based data expansion, we observed high reconstruction accuracy. We also observed low error with motion inference for rhythmic tasks, like treadmill walking or running. However, SHRED struggled to reliably and robustly infer motion during more complex tasks, like during environment exploration within the Total-Capture experiment. Visual inspection of the TotalCapture videos revealed high variability in how subjects explored their environment. Some of the tasks, such as acting or dancing, were often performed with little lower extremity movement and greater upper extremity movement, which may explain the inconsistent accuracy in data expansion for our focus on lower extremity motion. Given that generative models – such as SHRED’s encoder-decoder networks or the diffusion model used by DiffusionPoser – are learning the probability distribution of the training data and thus using the learned distribution to generate new, statistically-similar data, it is vital that we obtain and leverage relevant training data for robust and reliable data expansion. Indeed, large datasets that reflect an individual’s movements within their natural environment have been used to train motion inference algorithms [32], [83], [84]. For example, Geissinger and Asbeck introduced a dataset containing 40 hours of unscripted motion and trained machine learning models for motion inference [45]. However, there has not be an explicit quantification of motion complexity or its effect on motion inference accuracy. If we can quantify the relationship between motion complexity and inference, we may improve data collection design and model training protocols to better capture meaningful data for robust motion inference algorithms agnostic to environment or task.

## V. Conclusion

Shallow recurrent decoder networks can expand human motion data by robustly mapping a sparse set of sensor measurements to a dense set. This mapping may be used for both data reconstruction and inference and can be personalized or group-based. The time history of sensor measurements encodes important information about human motion and can be leveraged for training motion inference algorithms. However, understanding the effect of motion complexity on inference accuracy must be further studied, to ensure robust and reliable inference algorithms, agnostic to environment or task. By using the methods proposed here to expand datasets, wearable sensors could be used for more continuous and less intrusive monitoring of human motion in natural environments. Expanding our datasets to unmeasured or unavailable quantities can impact clinical trials, robotic/device control, and human performance by improving the accuracy and availability of digital biomarker estimates.

## Acknowledgment

The authors acknowledge support by the US National Science Foundation (NSF) AI Institute for Dynamical Systems (dynamicsai.org) grant 2112085. The authors also acknowledge funding in part by the National Institutes of Health (NIH) National Institute of Neurological Disorders and Stroke award R01NS091056.

## References

[1] J. P. Williams, O. Zahn, and J. N. Kutz, “Sensing with shallow recurrent decoder networks,” arXiv preprint 2301.12011, 2023.

[2] M. R. Ebers, J. P. Williams, K. M. Steele, and J. N. Kutz, “Leveraging arbitrary mobile sensor trajectories with shallow recurrent decoder networks for full-state reconstruction,” arXiv preprint 2307.11793, 2023.

[3] V. T. Inman, H. J. Ralston, and F. Todd, Human walking. Williams & Wilkins, 1981.

[4] B. van Veen, E. Montefiori, L. Modenese, C. Mazzà, and M. Viceconti, “Muscle recruitment strategies can reduce joint loading during level walking,” Journal of biomechanics, vol. 97, p. 109368, 2019.

[5] C. A. Myers, P. J. Laz, K. B. Shelburne, D. L. Judd, J. D. Winters, J. E. Stevens-Lapsley, and B. S. Davidson, “Simulated hip abductor strengthening reduces peak joint contact forces in patients with total hip arthroplasty,” Journal of biomechanics, vol. 93, pp. 18–27, 2019.

[6] M. J. Decker, M. R. Torry, T. J. Noonan, W. I. Sterett, and J. R. Steadman, “Gait retraining after anterior cruciate ligament reconstruction,” Archives of physical medicine and rehabilitation, vol. 85, no. 5, pp. 848–856, 2004.

[7] J. R. Rebula, L. V. Ojeda, P. G. Adamczyk, and A. D. Kuo, “Measurement of foot placement and its variability with inertial sensors,” Gait & posture, vol. 38, no. 4, pp. 974–980, 2013.

[8] T. E. Hewett, G. D. Myer, K. R. Ford, R. S. Heidt Jr, A. J. Colosimo, S. G. McLean, A. J. Van den Bogert, M. V. Paterno, and P. Succop, “Biomechanical measures of neuromuscular control and valgus loading of the knee predict anterior cruciate ligament injury risk in female athletes: a prospective study,” The American journal of sports medicine, vol. 33, no. 4, pp. 492–501, 2005.

[9] C. A. Okoro, “Prevalence of disabilities and health care access by disability status and type among adults—united states, 2016,” MMWR. Morbidity and mortality weekly report, vol. 67, 2018.

[10] A. Cieza, K. Causey, K. Kamenov, S. W. Hanson, S. Chatterji, and T. Vos, “Global estimates of the need for rehabilitation based on the global burden of disease study 2019: a systematic analysis for the global burden of disease study 2019,” The Lancet, vol. 396, no. 10267, pp. 2006–2017, 2020.

[11] P. Picerno, “25 years of lower limb joint kinematics by using inertial and magnetic sensors: A review of methodological approaches,” Gait & posture, vol. 51, pp. 239–246, 2017.

[12] E. Ceseracciu, Z. Sawacha, and C. Cobelli, “Comparison of markerless and marker-based motion capture technologies through simultaneous data collection during gait: proof of concept,” PloS one, vol. 9, no. 3, p. e87640, 2014.

[13] D. Hartley, “Rural health disparities, population health, and rural culture,” American journal of public health, vol. 94, no. 10, pp. 1675–1678, 2004.

[14] P. Braveman, “Health disparities and health equity: concepts and measurement,” Annu. Rev. Public Health, vol. 27, pp. 167–194, 2006.

[15] X. Chen, “Human motion analysis with wearable inertial sensors,” 2013.

[16] T. Cudejko, K. Button, J. Willott, and M. Al-Amri, “Applications of wearable technology in a real-life setting in people with knee osteoarthritis: A systematic scoping review,” Journal of Clinical Medicine, vol. 10, no. 23, p. 5645, 2021.

[17] T. McGrath and L. Stirling, “Body-worn imu-based human hip and knee kinematics estimation during treadmill walking,” Sensors, vol. 22, no. 7, p. 2544, 2022.

[18] S. Z. Homayounfar and T. L. Andrew, “Wearable sensors for monitoring human motion: a review on mechanisms, materials, and challenges,” SLAS TECHNOLOGY: Translating Life Sciences Innovation, vol. 25, no. 1, pp. 9–24, 2020.

[19] I. H. Lopez-Nava and A. Munoz-Melendez, “Wearable inertial sensors for human motion analysis: A review,” IEEE Sensors Journal, vol. 16, no. 22, pp. 7821–7834, 2016.

[20] W. De Vries, H. Veeger, C. Baten, and F. Van Der Helm, “Magnetic distortion in motion labs, implications for validating inertial magnetic sensors,” Gait & posture, vol. 29, no. 4, pp. 535–541, 2009.

[21] S. Zhang, Y. Li, S. Zhang, F. Shahabi, S. Xia, Y. Deng, and N. Alshurafa, “Deep learning in human activity recognition with wearable sensors: A review on advances,” Sensors, vol. 22, no. 4, p. 1476, 2022.

[22] F. Attal, S. Mohammed, M. Dedabrishvili, F. Chamroukhi, L. Oukhellou, and Y. Amirat, “Physical human activity recognition using wearable sensors,” Sensors, vol. 15, no. 12, pp. 31 314–31 338, 2015.

[23] S. Ramasamy Ramamurthy and N. Roy, “Recent trends in machine learning for human activity recognition—a survey,” Wiley Interdisciplinary Reviews: Data Mining and Knowledge Discovery, vol. 8, no. 4, p. e1254, 2018.

[24] G. M. Weiss, J. L. Timko, C. M. Gallagher, K. Yoneda, and A. J. Schreiber, “Smartwatch-based activity recognition: A machine learning approach,” in 2016 IEEE-EMBS International Conference on Biomedical and Health Informatics (BHI). IEEE, 2016, pp. 426–429.

[25] H. F. Nweke, Y. W. Teh, M. A. Al-Garadi, and U. R. Alo, “Deep learning algorithms for human activity recognition using mobile and wearable sensor networks: State of the art and research challenges,” Expert Systems with Applications, vol. 105, pp. 233–261, 2018.

[26] D. Biswas, A. Cranny, N. Gupta, K. Maharatna, J. Achner, J. Klemke, M. Jöbges, and S. Ortmann, “Recognizing upper limb movements with wrist worn inertial sensors using k-means clustering classification,” Human movement science, vol. 40, pp. 59–76, 2015.

[27] Z. Cao, T. Simon, S.-E. Wei, and Y. Sheikh, “Realtime multi-person 2d pose estimation using part affinity fields,” in Proceedings of the IEEE conference on computer vision and pattern recognition, 2017, pp. 7291–7299.

[28] H.-S. Fang, J. Li, H. Tang, C. Xu, H. Zhu, Y. Xiu, Y.-L. Li, and C. Lu, “Alphapose: Whole-body regional multi-person pose estimation and tracking in real-time,” IEEE Transactions on Pattern Analysis and Machine Intelligence, 2022.

[29] A. Mathis, P. Mamidanna, K. M. Cury, T. Abe, V. N. Murthy, M. W. Mathis, and M. Bethge, “Deeplabcut: markerless pose estimation of user-defined body parts with deep learning,” Nature neuroscience, vol. 21, no. 9, pp. 1281–1289, 2018.

[30] P. M. S. Ribeiro, A. C. Matos, P. H. Santos, and J. S. Cardoso, “Machine learning improvements to human motion tracking with imus,” Sensors, vol. 20, no. 21, p. 6383, 2020.

[31] T. Von Marcard, B. Rosenhahn, M. J. Black, and G. Pons-Moll, “Sparse inertial poser: Automatic 3d human pose estimation from sparse imus,” in Computer graphics forum, vol. 36, o. 2. Wiley Online Library, 2017, pp. 349–360.

[32] Y. Huang, M. Kaufmann, E. Aksan, M. J. Black, O. Hilliges, and G. Pons-Moll, “Deep inertial poser: Learning to reconstruct human pose from sparse inertial measurements in real time,” ACM Transactions on Graphics (TOG), vol. 37, no. 6, pp. 1–15, 2018.

[33] L. A. Schwarz, D. Mateus, and N. Navab, “Discriminative human full-body pose estimation from wearable inertial sensor data,” in Modelling the Physiological Human: 3D Physiological Human Workshop, 3DPH 2009, Zermatt, Switzerland, November 29–December 2, 2009. Proceedings. Springer, 2009, pp. 159–172.

[34] F. J. Wouda, M. Giuberti, G. Bellusci, and P. H. Veltink, “Estimation of full-body poses using only five inertial sensors: an eager or lazy learning approach?” Sensors, vol. 16, no. 12, p. 2138, 2016.

[35] M. Luštrek and B. Kaluža, “Fall detection and activity recognition with machine learning,” Informatica, vol. 33, no. 2, 2009.

[36] G. L. Santos, P. T. Endo, K. H. d. C. Monteiro, E. d. S. Rocha Silva, and T. Lynn, “Accelerometer-based human fall detection using convolutional neural networks,” Sensors, vol. 19, no. 7, p. 1644, 2019.

[37] F. Hussain, F. Hussain, M. Ehatisham-ul Haq, and M. A. Azam, “Activity-aware fall detection and recognition based on wearable sensors,” IEEE Sensors Journal, vol. 19, no. 12, pp. 4528–4536, 2019.

[38] T.-S. Wei, P.-T. Liu, L.-W. Chang, and S.-Y. Liu, “Gait asymmetry, ankle spasticity, and depression as independent predictors of falls in ambulatory stroke patients,” PloS one, vol. 12, no. 5, p. e0177136, 2017.

[39] D. Roetenberg, P. J. Slycke, and P. H. Veltink, “Ambulatory position and orientation tracking fusing magnetic and inertial sensing,” IEEE Transactions on Biomedical Engineering, vol. 54, no. 5, pp. 883–890, 2007.

[40] E. Rapp, S. Shin, W. Thomsen, R. Ferber, and E. Halilaj, “Estimation of kinematics from inertial measurement units using a combined deep learning and optimization framework,” Journal of Biomechanics, vol. 116, p. 110229, 2021.

[41] P. Tsinganos and A. Skodras, “On the comparison of wearable sensor data fusion to a single sensor machine learning technique in fall detection,” Sensors, vol. 18, no. 2, p. 592, 2018.

[42] H. F. Nweke, Y. W. Teh, U. R. Alo, and G. Mujtaba, “Analysis of multi-sensor fusion for mobile and wearable sensor based human activity recognition,” in Proceedings of the international conference on data processing and applications, 2018, pp. 22–26.

[43] R. C. King, E. Villeneuve, R. J. White, R. S. Sherratt, W. Holderbaum, and W. S. Harwin, “Application of data fusion techniques and technologies for wearable health monitoring,” Medical engineering & physics, vol. 42, pp. 1–12, 2017.

[44] D. Vlasic, R. Adelsberger, G. Vannucci, J. Barnwell, M. Gross, W. Matusik, and J. Popović, “Practical motion capture in everyday surroundings,” ACM transactions on graphics (TOG), vol. 26, no. 3, pp. 35–es, 2007.

[45] J. H. Geissinger and A. T. Asbeck, “Motion inference using sparse inertial sensors, self-supervised learning, and a new dataset of unscripted human motion,” Sensors, vol. 20, no. 21, p. 6330, 2020.

[46] A. Findlow, J. Goulermas, C. Nester, D. Howard, and L. Kenney, “Predicting lower limb joint kinematics using wearable motion sensors,” Gait & posture, vol. 28, no. 1, pp. 120–126, 2008.

[47] J. L. McGinley, R. Baker, R. Wolfe, and M. E. Morris, “The reliability of three-dimensional kinematic gait measurements: a systematic review,” Gait & posture, vol. 29, no. 3, pp. 360–369, 2009.

[48] K. Manohar, B. W. Brunton, J. N. Kutz, and S. L. Brunton, “Data-driven sparse sensor placement for reconstruction: Demonstrating the benefits of exploiting known patterns,” IEEE Control Systems Magazine, vol. 38, no. 3, pp. 63–86, 2018.

[49] T. Bolton and L. Zanna, “Applications of deep learning to ocean data inference and subgrid parameterization,” Journal of Advances in Modeling Earth Systems, vol. 11, no. 1, pp. 376–399, 2019.

[50] J. L. Callaham, K. Maeda, and S. L. Brunton, “Robust flow reconstruction from limited measurements via sparse representation,” Physical Review Fluids, vol. 4, no. 10, p. 103907, 2019.

[51] J. Yu and J. S. Hesthaven, “Flowfield reconstruction method using artificial neural network,” Aiaa Journal, vol. 57, no. 2, pp. 482–498, 2019.

[52] N. B. Erichson, L. Mathelin, Z. Yao, S. L. Brunton, M. W. Mahoney, and J. N. Kutz, “Shallow neural networks for fluid flow reconstruction with limited sensors,” Proceedings of the Royal Society A, vol. 476, no. 2238, p. 20200097, 2020.

[53] D. P. Kingma and J. Ba, “Adam: A method for stochastic optimization,” arXiv preprint 1412.6980, 2014.

[54] M. C. Rosenberg, B. S. Banjanin, S. A. Burden, and K. M. Steele, “Predicting walking response to ankle exoskeletons using data-driven models,” Journal of the Royal Society Interface, vol. 17, no. 171, p. 20200487, 2020.

[55] A. Rajagopal, Ł. Kidziński, A. S. McGlaughlin, J. L. Hicks, S. L. Delp, and M. H. Schwartz, “Estimating the effect size of surgery to improve walking in children with cerebral palsy from retrospective observational clinical data,” Scientific reports, vol. 8, no. 1, pp. 1–11, 2018.

[56] J. L. Hicks, T. K. Uchida, A. Seth, A. Rajagopal, and S. L. Delp, “Is my model good enough? best practices for verification and validation of musculoskeletal models and simulations of movement,” Journal of biomechanical engineering, vol. 137, no. 2, 2015.

[57] G. Rau, C. Disselhorst-Klug, and R. Schmidt, “Movement biomechanics goes upwards: from the leg to the arm,” Journal of biomechanics, vol. 33, no. 10, pp. 1207–1216, 2000.

[58] T.-W. Lu and C.-F. Chang, “Biomechanics of human movement and its clinical applications,” The Kaohsiung journal of medical sciences, vol. 28, pp. S13–S25, 2012.

[59] K. A. Ingraham, D. P. Ferris, and C. D. Remy, “Evaluating physiological signal salience for estimating metabolic energy cost from wearable sensors,” Journal of applied physiology, vol. 126, no. 3, pp. 717–729, 2019.

[60] L. C. Benson, A. M. Räisänen, C. A. Clermont, and R. Ferber, “Is this the real life, or is this just laboratory? a scoping review of imu-based running gait analysis,” Sensors, vol. 22, no. 5, p. 1722, 2022.

[61] A. M. Spomer, R. Z. Yan, M. H. Schwartz, and K. M. Steele, “Motor control complexity can be dynamically simplified during gait pattern exploration using motor control-based biofeedback,” Journal of neurophysiology, vol. 129, no. 5, pp. 984–998, 2023.

[62] K. Pearson, “Liii. on lines and planes of closest fit to systems of points in space,” The London, Edinburgh, and Dublin philosophical magazine and journal of science, vol. 2, no. 11, pp. 559–572, 1901.

[63] S. Wold, K. Esbensen, and P. Geladi, “Principal component analysis,” Chemometrics and intelligent laboratory systems, vol. 2, no. 1-3, pp. 37–52, 1987.

[64] H. Abdi and L. J. Williams, “Principal component analysis,” Wiley interdisciplinary reviews: computational statistics, vol. 2, no. 4, pp. 433–459, 2010.

[65] M. Trumble, A. Gilbert, C. Malleson, A. Hilton, and J. Collomosse, “Total capture: 3d human pose estimation fusing video and inertial sensors,” in 2017 British Machine Vision Conference (BMVC), 2017.

[66] T. Van Wouwe, S. Lee, A. Falisse, S. Delp, and C. K. Liu, “Diffusion inertial poser: Human motion reconstruction from arbitrary sparse imu configurations,” arXiv preprint 2308.16682, 2023.

[67] K. Werling, N. A. Bianco, M. Raitor, J. Stingel, J. L. Hicks, S. H. Collins, S. L. Delp, and C. K. Liu, “Addbiomechanics: Automating model scaling, inverse kinematics, and inverse dynamics from human motion data through sequential optimization,” Plos one, vol. 18, no. 11, p. e0295152, 2023.

[68] N. Takayanagi, M. Sudo, Y. Yamashiro, S. Lee, Y. Kobayashi, Y. Niki, and H. Shimada, “Relationship between daily and in-laboratory gait speed among healthy community-dwelling older adults,” Scientific reports, vol. 9, no. 1, p. 3496, 2019.

[69] S. Del Din, A. Godfrey, B. Galna, S. Lord, and L. Rochester, “Freeliving gait characteristics in ageing and parkinson’s disease: impact of environment and ambulatory bout length,” Journal of neuroengineering and rehabilitation, vol. 13, pp. 1–12, 2016.

[70] S. Shema-Shiratzky, I. Hillel, A. Mirelman, K. Regev, K. L. Hsieh, A. Karni, H. Devos, J. J. Sosnoff, and J. M. Hausdorff, “A wearable sensor identifies alterations in community ambulation in multiple sclerosis: contributors to real-world gait quality and physical activity,” Journal of neurology, vol. 267, pp. 1912–1921, 2020.

[71] R. D. Gurchiek, R. H. Choquette, B. D. Beynnon, J. R. Slauterbeck, T. W. Tourville, M. J. Toth, and R. S. McGinnis, “Remote gait analysis using wearable sensors detects asymmetric gait patterns in patients recovering from acl reconstruction,” in 2019 IEEE 16th International Conference on Wearable and Implantable Body Sensor Networks (BSN). IEEE, 2019, pp. 1–4.

[72] R. E. Van Emmerik, S. W. Ducharme, A. C. Amado, and J. Hamill, “Comparing dynamical systems concepts and techniques for biomechanical analysis,” Journal of sport and health science, vol. 5, no. 1, pp. 3–13, 2016.

[73] G. P. Austin, “Motor control of human gait: A dynamic systems perspective,” 2001.

[74] J. Hamill, R. E. van Emmerik, B. C. Heiderscheit, and L. Li, “A dynamical systems approach to lower extremity running injuries,” Clinical biomechanics, vol. 14, no. 5, pp. 297–308, 1999.

[75] C. Basdogan and F. M. Amirouche, “Nonlinear dynamics of human locomotion: from the perspective of dynamical systems theory,” in Engineering Systems Design and Analysis Conference, 1996.

[76] L. Pitto, H. Kainz, A. Falisse, M. Wesseling, S. Van Rossom, H. Hoang, E. Papageorgiou, A. Hallemans, K. Desloovere, G. Molenaers et al., “Simcp: A simulation platform to predict gait performance following orthopedic intervention in children with cerebral palsy,” Frontiers in neurorobotics, vol. 13, p. 54, 2019.

[77] A. S. Arnold, S. S. Blemker, and S. L. Delp, “Evaluation of a deformable musculoskeletal model for estimating muscle–tendon lengths during crouch gait,” Annals of biomedical engineering, vol. 29, pp. 263–274, 2001.

[78] L. Scheys, A. Van Campenhout, A. Spaepen, P. Suetens, and I. Jonkers, “Personalized mr-based musculoskeletal models compared to rescaled generic models in the presence of increased femoral anteversion: effect on hip moment arm lengths,” Gait & posture, vol. 28, no. 3, pp. 358–365, 2008.

[79] P. Hartman, “Ordinary differential equations, classics in applied mathematics, vol. 38 (society for industrial and applied mathematics (siam), philadelphia, pa),” corrected reprint of the second (1982) edition [Birkhäuser, Boston, MA, 2002.

[80] S. M. Hirsh, S. M. Ichinaga, S. L. Brunton, J. Nathan Kutz, and B. W. Brunton, “Structured time-delay models for dynamical systems with connections to frenet–serret frame,” Proceedings of the Royal Society A, vol. 477, no. 2254, p. 20210097, 2021.

[81] J. Bakarji, K. Champion, J. N. Kutz, and S. L. Brunton, “Discovering governing equations from partial measurements with deep delay autoen-coders,” arXiv preprint 2201.05136, 2022.

[82] F. Takens, “Detecting strange attractors in turbulence,” in Dynamical Systems and Turbulence, Warwick 1980: proceedings of a symposium held at the University of Warwick 1979/80. Springer, 2006, pp. 366–381.

[83] N. Mahmood, N. Ghorbani, N. F. Troje, G. Pons-Moll, and M. J. Black, “Amass: Archive of motion capture as surface shapes,” in Proceedings of the IEEE/CVF international conference on computer vision, 2019, pp. 5442–5451.

[84] V. Mollyn, R. Arakawa, M. Goel, C. Harrison, and K. Ahuja, “Imuposer: Full-body pose estimation using imus in phones, watches, and earbuds,” in Proceedings of the 2023 CHI Conference on Human Factors in Computing Systems, 2023, pp. 1–12.

